## Supplemental data for "Organising the cell cycle in the absence of transcriptional control: Dynamic phosphorylation co-ordinates the *Trypanosoma brucei* cell cycle post-transcriptionally"

#### Supplementary Data

Figure S9. Venn diagram of the overlap of the two CCR proteomes and the CCR transcriptome. .. 14

#### Additional Supplementary Tables *(separate excel spreadsheets)*

**Table S2.** 5,949 Phosphorylation sites quantified at all six time points.

**Table S3.** 3,619 Proteins quantified at all six time points.

**Table S4.** 917 cell cycle regulated phosphorylation sites.

**Table S5.** 443 Cell cycle regulated proteins.

**Table S6**. Immunoprecipitation of *T. brucei* CSBPII

#### Supplementary Materials & Methods

Cell culture and SILAC labelling

The Stable isotope labelling by amino acids in cell culture (SILAC) labelling of *T. brucei* 427 Lister procyclic form cells was performed as described previously (18, 20). In brief, log phase parasites were passaged into SDM-79 SILAC media (SDM-79 medium lacking L-arginine and L-lysine) supplemented with 10% dialysed FCS (1000 MWCO, Dundee Cell Products) and the standard concentration of either normal isotopic abundance L-arginine and L-lysine (SDM-79, referred to as Light label) or $^{13}C_6$ L-arginine and 4,4,5,5-$D_4$ L-lysine (SDM-79 + R6K4, referred to as Medium label) or with $^{13}C_6$,$^{15}N_4$L-arginine and $^{13}C_6$,$^{15}N_2$ L-lysine (SDM-79 + R10K8, referred to as Heavy label). The stable isotope-labelled amino acids were obtained from CK Isotopes, UK. Cells were allowed to grow in the respective medium for more than 7 cell divisions before cultures were enlarged to a volume of 200-300 ml at a final concentration of 1-2 × $10^7$ cells/ml for the elutriated samples (light and heavy label). To produce the medium-SILAC standard for use as an asynchronous control, a single batch of 1L of medium-labelled cells was grown to a density of 1 × $10^7$ cells/ml to minimise variability.

Synchronisation by centrifugal counter flow elutriation

Elutriation was performed essentially as described in (19). Briefly, logarithmic phase Pcf cells (5 × $10^6$ - 1.5 × $10^7$/ml) were harvested by centrifugation at 1,000 × g for 10 min at room temperature (RT) and resuspended in 20 ml Pcf elutriation buffer (PBS + 25% SDM-79). A maximum number of 3 × $10^9$ Pcf were used for each elutriation. For early and late G1 samples, 150 ml fractions eluting at 18 and 20 ml/min (early) and 22 and 24 ml/min (late) were harvested by centrifugation and processed for proteomic analysis as described below. For some late G2/M samples, the pump speed was gradually increased and fractions of about 100 ml collected and discarded to deplete the sample of G1 and S phase cells. The remaining cells (highly enriched in G2/M) were eventually collected at a pump speed of 35 ml/min. For any other samples, 150 ml fractions eluting at 18, 20 and 22 ml/min were harvested by centrifugation and resuspended in the appropriate amount of SDM-79 for further cultivation. Samples for proteomics and flow cytometry were withdrawn at appropriate time points.

To follow up on specific proteins in cell lines bearing epitope-tagged versions of the respective proteins, elutriation was performed as described above except with slightly smaller culture sizes (100-150ml). For SmOxP9-based cell lines (51), 100 ml fractions were collected and the fraction eluting at 18 ml/min was harvested by centrifugation and resuspended in the appropriate amount of SDM-79 for further cultivation. Flow cytometry samples, cell lysates for western blots and microscopy slides were prepared at different time points following elutriation as specified in the respective figures.

Filter aided sample preparation (FASP)

Synchronised samples from two time points (light- and heavy-labelled) and one asynchronous control sample (medium-labelled) were thawed and mixed (2 × $10^8$ cells per sample, giving a total of 6 × $10^8$ cells). The proteins were solubilized with SDS and tryptic peptides generated by an adaptation of the filter aided sample preparation procedure as described previously (18, 22), except using a 1:50 w/w trypsin to protein ration of Trypsin Gold (Promega). The digested peptide solution was then removed from the column, diluted to 3 ml with ABC and acidified with 0.1% TFA before desalting using a 500 mg $C_{18}$ column (SepPak, Waters), lyophilisation and storage at -80 $^0$C.

Separation of peptides from phosphopeptides by Fe-IMAC

Separation of peptides from phosphopeptides using Fe-IMAC was performed as described by Ruprecht *et al* (23) with minor modifications. An analytical Fe-IMAC column (4 × 50 mm ProPac IMAC-10, Thermo Fisher Scientific) connected to an LC Packing Famos HPLC was charged with $FeCl_3$ as described in (Ruprecht et al. 2015). Buffer A (30% MeCN, 0.07% TFA) was then used to wash and equilibrate the column before a blank run was performed. The gradient consisted of 0-3 min 100% buffer A at 50 µl/min, 3-4 min 100% buffer A at 50-150 µl/min, 4-15 min 100% buffer A at 150 µl/min, 15-16 min 100% buffer A at 150-100 µl/min, 16-76min 0-45% buffer B (30% MeCN, 0.5% $NH_4OH$) at 100 µl/min, 76-77 min 100% buffer B at 100 µl/min, 77-82 min 100% buffer B at 100-200 µl/min, 82-85 min 100-0% buffer B at 200µl/min and 85-115min 100% buffer A at 200 µl/min. Lyophilised tryptic digests were resuspended in 100 µl of buffer A before injection onto the column. Fractions were collected every 2 min in low protein binding micro centrifuge tubes (Eppendorf Protein lo-bind), with elution of peptides monitored by absorbance at 280 nm. Early eluting fractions (<6 min) corresponding to peptides and later eluting fractions corresponding to phosphopeptides (~30 min) were lyophilised and stored at -80 $^0$C prior to further processing.

<u>High pH reverse phase fractionation</u>

Samples were fractionated using the High pH Fractionation kit (ThermoFisher) according to the manufacturer's instructions with the following modifications. Lyophilised (phospho)peptides were resuspended in 100µl 5 mM $NH_4OH$ and applied to the pre-treated columns (MeCN, 0.1 % TFA, and two 5 mM $NH_4OH$ washes). The flow-through was reapplied before being collected and desalted using $C_{18}$ microspin columns (Harvard Apparatus). The columns were then washed 3 times with 5 mM $NH_4OH$ and the (phospho)peptides eluted with different concentrations of MeCN (2, 3, 4, 6, 10 and 50% MeCN). The eluates were concatenated into five fractions as follows: F1 = 2% + 50% eluates, F2 = 3% eluate, F3 = 4% eluate, F4 = 6% eluate, F5 = 10% eluate + desalted flow through. The concatenated fractions were then lyophilised prior to analysis by LC-MS/MS.

<u>Selection of cell cycle regulated proteins and phosphorylation sites</u>

Prior to further analysis, the outputs from MaxQuant (4) were filtered to remove known contaminants and reverse sequences, and any SILAC ratios with greater ≥100% variation or phosphorylation sites with a localisation probability < 0.95 were excluded. The 18 biological replicates were grouped into six cell cycle time points (early G1, EG1; late G1, LG1; early S, ES; late S, LS, early G2/M, EG2M; late G2M, LG2M) on the basis of the proportion of cells in G1, S and G2/M phases determined by flow cytometry analysis of propidium iodide stained cells (Supplemental Table S1). The SILAC ratios of the individual time points were averaged, and the data set filtered to remove any protein groups or phosphorylation sites that were not observed at all six cell cycle time points. The averaged ratios obtained compare an individual cell cycle time point (i.e. EG1, LS) to the same medium labelled asynchronous population ($M_{Asyn}$), which therefore constitutes an invariable internal standard that can be used to directly compare different cell cycle time points. We used the averaged EG1 sample (95-97% G1), which represents the start of the cell cycle, to normalize the data to give ratios relative to the abundance at the start of the cell cycle.

$$Log_2 \frac{LS}{EG1} = Log_2 \frac{L_{LS}}{M_{Asyn}} - Log_2 \frac{H_{EG1}}{M_{Asyn}}$$

The maximum fold change ($FC_{Max}$) across the cell cycle was calculated as the difference between the maximum and minimum abundance relative to EG1, and any protein with an $FC_{Max}$ ≥ 1.5-fold or phosphorylation site with $FC_{Max}$ ≥ 3-fold was considered to be cell cycle regulated (CCR). To make our data accessible to the scientific community, we have submitted the results of our study to TriTrypDB, enabling researchers to access the data presented here.

<u>Peak time calculation</u>

To determine the order in which the CCR protein/phosphorylation site abundances reaches a maximum, peak time $t_{peak}$ was calculated according to the procedure of Olsen et al (5). The ratios (averaged, normalised to EG1) $r_1$ to $r_6$ were scaled to the unit interval [0, 1] and the six time points $t_1$

to $t_6$ assigned values 1 to 6. Then for each protein the peak time $t_{peak}$ was calculated using a weighted mean of the expression ratio of maximal expression ($r_i = 1$) at time point $t_i$ with respect to the adjacent time points ($t_{i-1}$ and $t_{i+1}$). Since the time points represent a continuous cycle, if the maximal expression was at $t_1$ it was preceded by $t_0$ with expression $r_6$ and if maximal expression was at $t_6$ it was followed by $t_7 = 7$ with expression $r_1$.

$$t_{peak} = \begin{cases} \dfrac{t_{i-1} \times r_{i-1} + t_i r_i + t_{i+1} \times r_{i+1}}{r_{i-1} + r_i + r_{i+1}}, \text{if} \max r_i \text{ at } i \in [2,5] \\ \dfrac{t_i r_i + t_{i+1} \times r_{i+1} + 0 \times r_6}{r_i + r_{i+1} + r_6}, \text{if} \max r_i \text{ at } i = 1 \\ \dfrac{t_{i-1} \times r_{i-1} + t_i r_i + 7 r_{i+1}}{r_{i-1} + r_i + r_7}, \text{if} \max r_i \text{ at } i = 6 \end{cases}$$

The profiles were then ordered in increasing $t_{peak}$ to gain temporal map of the cell cycle rendered as a heat map of the unit interval scaled values, and projected on a polar plot of $t_{peak}$ versus $FC_{Max}$.

Hierarchical clustering

To cluster the regulated protein/phosphorylation sites that show similar expression profiles, the ratios in each expression profile were Z-scored by subtracting the mean profile value and dividing by the standard deviation, and hierarchical clustering performed using Euclidean distance of the complete linkage after pre-processing with K-means using 1000 iteration and 10 restarts of 300 or 150 clusters for phosphorylation sites or proteins respectively (6). The final clusters were defined using a minimum distance threshold of < 2.5, resulting in 30 phosphorylation site clusters and 29 protein abundance clusters.

Cluster annotation enrichment analysis

Gene ontology (GO) term enrichment analysis of clusters was performed using the TriTrypDB curated GO Slim set contained within the integrated GO enrichment tool , using an uncorrected *P* value cut off of 0.05 (7). Kinase motif enrichment analysis was conducted in Perseus (6) by searching the protein sequence flanking the identified phosphorylation site for linear motifs corresponding to potential kinase motifs, and using Fisher's exact test to determine the correlation between the phosphorylation site clusters and identified kinase motifs using a threshold of a Benjamini-Hochberg false discovery rate < 0.05. In addition, previously identified PLK substrates and binders (8, 9) were mapped onto phosphorylation sites at the protein level, and enrichment examined using Fisher's exact test as above.

### References

1. Parsons M, Worthey EA, Ward PN, Mottram JC. Comparative analysis of the kinomes of three pathogenic trypanosomatids: Leishmania major, Trypanosoma brucei and Trypanosoma cruzi. BMC Genomics. 2005;6:127.
2. Lueong S, Merce C, Fischer B, Hoheisel JD, Erben ED. Gene expression regulatory networks in Trypanosoma brucei: insights into the role of the mRNA-binding proteome. Mol Microbiol. 2016;100(3):457-71.
3. Erben ED, Fadda A, Lueong S, Hoheisel JD, Clayton C. A genome-wide tethering screen reveals novel potential post-transcriptional regulators in Trypanosoma brucei. PLoS Pathog. 2014;10(6):e1004178.
4. Cox J, Mann M. MaxQuant enables high peptide identification rates, individualized p.p.b.-range mass accuracies and proteome-wide protein quantification. Nat Biotechnol. 2008;26(12):1367-72.

5. Olsen JV, Vermeulen M, Santamaria A, Kumar C, Miller ML, Jensen LJ, et al. Quantitative phosphoproteomics reveals widespread full phosphorylation site occupancy during mitosis. Sci Signal. 2010;3(104):ra3.
6. Tyanova S, Temu T, Sinitcyn P, Carlson A, Hein MY, Geiger T, et al. The Perseus computational platform for comprehensive analysis of (prote)omics data. Nat Methods. 2016;13(9):731-40.
7. Aslett M, Aurrecoechea C, Berriman M, Brestelli J, Brunk BP, Carrington M, et al. TriTrypDB: a functional genomic resource for the Trypanosomatidae. Nucleic Acids Res. 2010;38(Database issue):D457-62.
8. McAllaster MR, Ikeda KN, Lozano-Nunez A, Anrather D, Unterwurzacher V, Gossenreiter T, et al. Proteomic identification of novel cytoskeletal proteins associated with TbPLK, an essential regulator of cell morphogenesis in Trypanosoma brucei. Mol Biol Cell. 2015;26(17):3013-29.
9. Hu H, Zhou Q, Li Z. A Novel Basal Body Protein That Is a Polo-like Kinase Substrate Is Required for Basal Body Segregation and Flagellum Adhesion in Trypanosoma brucei. J Biol Chem. 2015;290(41):25012-22.

#### Supplementary Tables & Figures

Table S1. Metadata for cell cycle proteomics samples.

| Sample Name | Sample origin[1] | Proteomic Experiment | G1[2] | S[2] | G2/M[2] |
|---|---|---|---|---|---|
| **EG1-a** | Direct, 18-20 ml/min | CC10-Heavy | 96 % | 3 % | 1 % |
| **EG1-b** | Direct, 18-20 ml/min | CC11-Heavy | 97 % | 2 % | 1 % |
| **EG1-c** | Direct, 18-20 ml/min | CC12-Light | 95 % | 2 % | 3 % |
| **LG1-a** | Direct, 22-24 ml/min | CC9-Heavy | 84 % | 14 % | 2 % |
| **LG1-b** | Direct, 22-24 ml/min | CC8-Heavy | 85 % | 12 % | 3 % |
| **LG1-c** | Direct, 22 ml/min | CC13-Light | 84 % | 11 % | 5 % |
| **ES-a** | Grown | CC14-Heavy | 45 % | 45 % | 10 % |
| **ES-b** | Grown | CC12-Heavy | 34 % | 45 % | 21 % |
| **ES-c** | Grown | CC6-Light | 40 % | 32 % | 28 % |
| **LS-a** | Grown | CC15-Heavy | 40 % | 48 % | 12 % |
| **LS-b** | Grown | CC14-Light | 22 % | 52 % | 26 % |
| **LS-c** | Grown | CC13-Heavy | 20 % | 46 % | 34 % |
| **EG2M-a** | Grown | CC9-Light | 16 % | 45 % | 39 % |
| **EG2M-b** | Grown | CC15-Light | 14 % | 47 % | 39 % |
| **EG2M-c** | Grown | CC8-Light | 19 % | 29 % | 52 % |
| **LG2M-a** | Grown | CC10-Light | 15 % | 25 % | 60 % |
| **LG2M-b** | Grown | CC11-Light | 24 % | 16 % | 60 % |
| **LG2M-c** | Direct, 25-35 ml/min | CC6-Heavy | 23 % | 12 % | 65 % |

[1] Elutriation conditions; either directly eluted or eluted and placed back into culture

[2] Cell cycle composition determined by flow cytometry of PI-stained sample.

Table S7. PSP1-C terminal domain containing proteins present in T. brucei. [1] RNA binding proteins that interact directly with mRNA (38, 39). NC, Not changing; -, not observed; +, RBP.

| Name | GeneDB ID | Proteome | Phosphorylation site | RBP[1] |
|---|---|---|---|---|
| **PCD1** | Tb927.11.14750 | LS | S159 | + |
| **PCD2** | Tb927.11.4180 | ES | S161, T132, S344 | + |
| **PCD3** | Tb927.10.8330 | ES | S81 | |
| **PCD4** | Tb927.10.9910 | NC | T100 | + |
| ***Tb*CSBPII-33** | Tb927.11.7140 | NC | S64, T68, S70, S85, S88 | + |
| ***Tb*CSBPII-45** | Tb927.5.760 | NC | S85 | |
| **PIE8** | Tb927.6.2850 | - | T144 | |
| **PCD5** | Tb927.10.11630 | LG2M | - | |
| **PCD6** | Tb927.8.3850 | - | - | + |
| **PCD7** | Tb927.3.5080 | - | - | |
| **PCD8** | Tb927.9.9370 | - | - | |

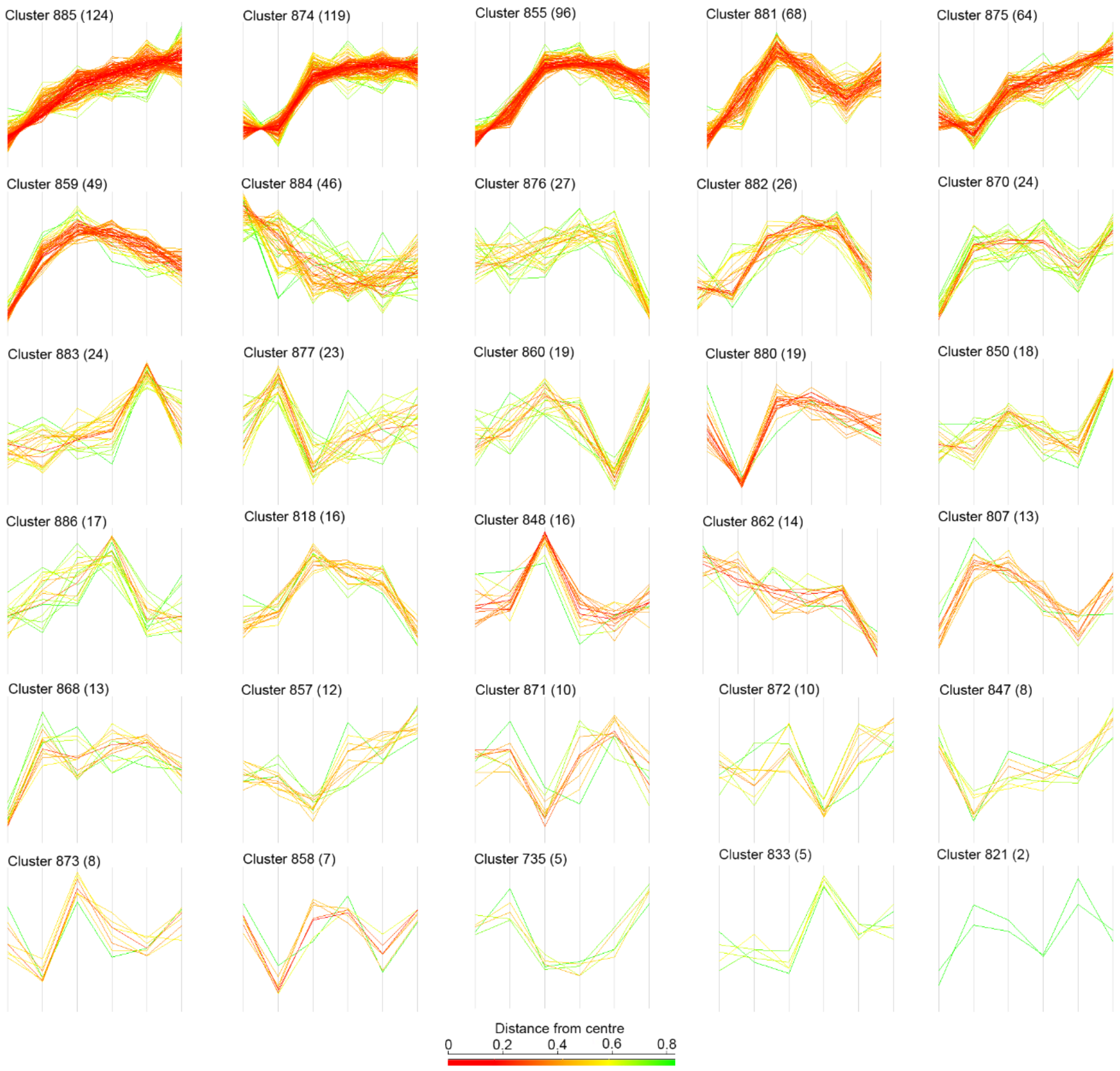

Figure S1. Hierarchical clustering of cell cycle regulated phosphorylation sites. The phosphorylation site ratios were Z-transformed and unrestrained hierarchical clustering performed using Euclidean distance of the complete linkage after pre-processing with K-means using 1000 iteration and 10 restarts of 300 clusters, with the final clusters were defined using a minimum distance threshold of < 2.5. The Cluster size is given in brackets, and the profiles are coloured by the Euclidean distance from the centre (mean profile) of the cluster.

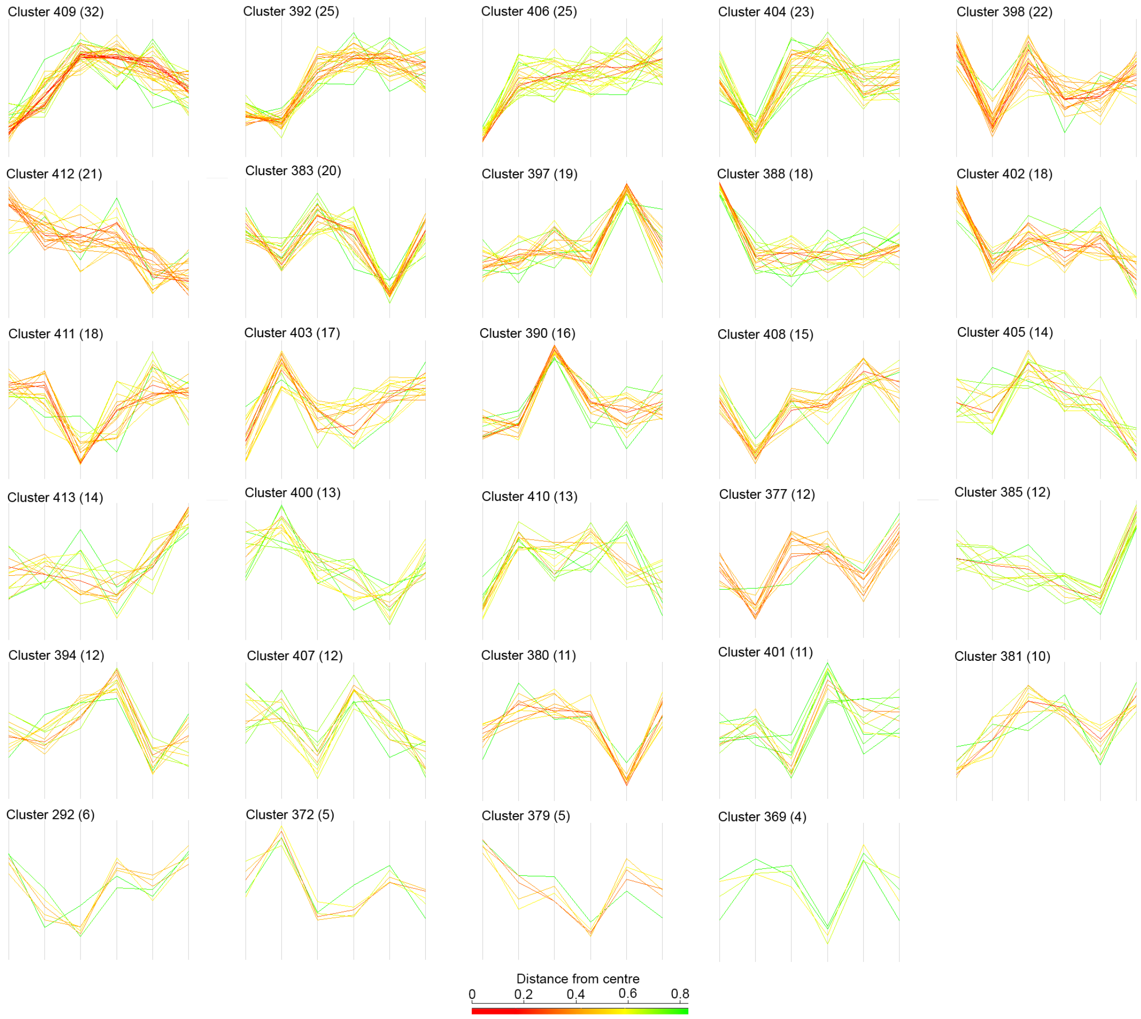

Figure S2. Hierarchical clustering of cell cycle regulated proteins. The protein ratios were Z-transformed and unrestrained hierarchical clustering performed using Euclidean distance of the complete linkage after pre-processing with K-means using 1000 iteration and 10 restarts of 150 clusters, with the final clusters were defined using a minimum distance threshold of < 2.5. The Cluster size is given in brackets, and the profiles are coloured by the Euclidean distance from the centre (mean profile) of the cluster.

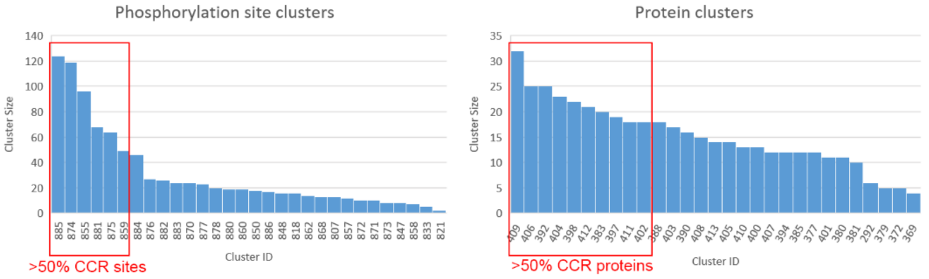

| GO term | 1 | 2 | 3 | 4 | 5 | 6 |
|---|---|---|---|---|---|---|
| Mitotic cell cycle | + | + | + | - | + | - |
| Cell cycle | + | + | + | - | - | + |
| Chromosome segregation | + | + | + | - | + | - |
| Cell division | + | + | - | - | - | + |
| mRNA binding | - | + | - | + | - | - |
| RNA binding | - | + | + | + | - | - |
| Chromosome | - | - | + | - | + | - |

| GO term | 1 | 2 | 3 | 4 | 5 | 6 | 7 | 8 | 9 | 10 |
|---|---|---|---|---|---|---|---|---|---|---|
| Mitotic cell cycle | + | - | + | - | - | - | + | - | - | - |
| Cell cycle | + | - | - | - | - | - | + | - | - | - |
| Chromosome segregation | - | + | - | - | - | - | - | - | - | - |
| Cell division | + | - | - | - | - | - | - | - | - | - |
| mRNA binding | - | - | + | - | - | - | - | - | - | - |
| RNA binding | - | - | - | - | - | - | - | - | - | - |
| Chromosome | - | + | + | - | - | - | - | - | - | - |

Figure S3. Comparison of Gene Ontology enrichment in cell cycle regulated phosphorylation site and protein clusters. Gene Ontology (GO) enrichment analysis using GO Slim ontology with $P < 0.05$ was performed on the largest clusters representing >50% of the CCR phosphorylation sites and proteins, and the occurrence of GO terms related to cell cycle tabulated.

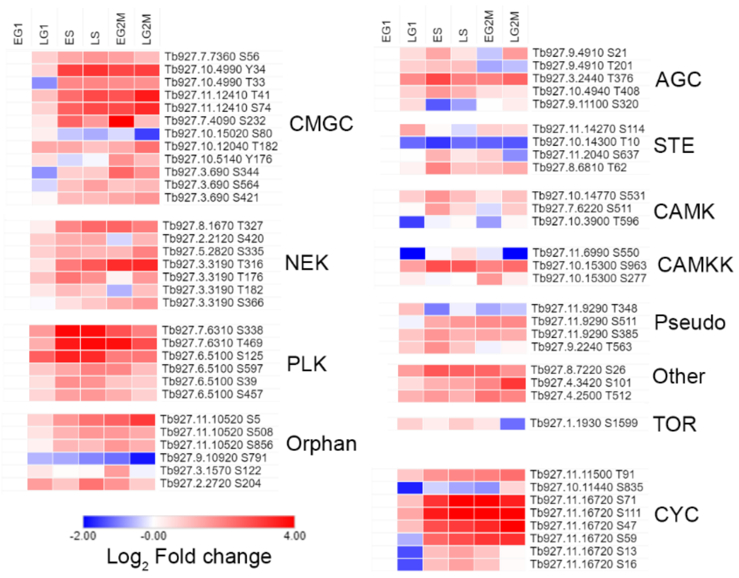

Figure S4. Heat map of cell cycle regulated protein kinase and cyclins. CCR protein kinase and cyclins rendered as a heat map of the $\log_2$ fold change relative to EG1, grouped by family (1).

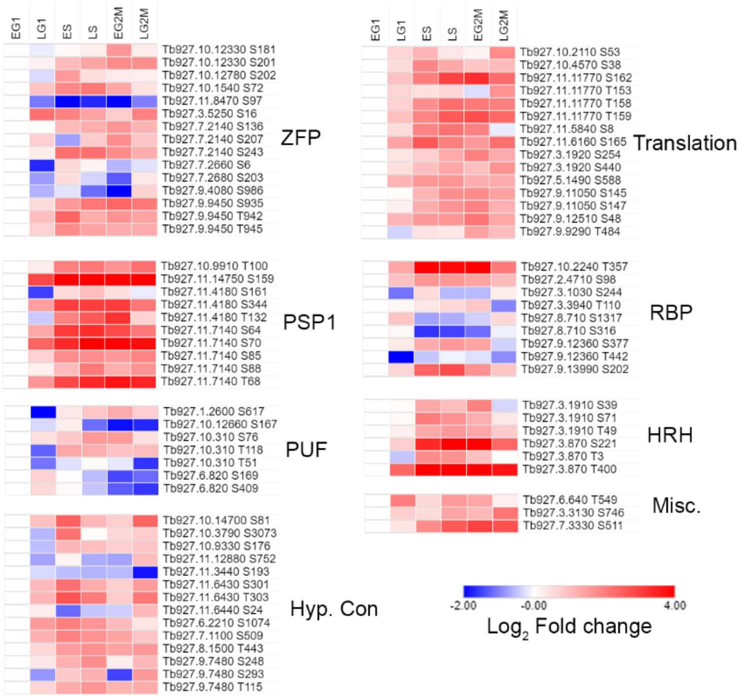

Figure S5. Heat map of cell cycle regulated RNA binding proteins. CCR Proteins containing recognisable RNA binding domains or identified from mRNA tethering screens and crosslinking proteomics (2, 3) are rendered as a heat map of the $\log_2$ fold change relative to EG1, grouped by proteins features. ZFP – zinc finger proteins; Translation – eIF and associated proteins; PSP1 – PSP1 C-terminal domain; RBP – RNA binding motif; PUF - Pumilio/Fem-3 domain; HRH – Histone RNA hairpin; Hyp. Con – hypothetical conserved proteins; Misc. – miscellaneous.

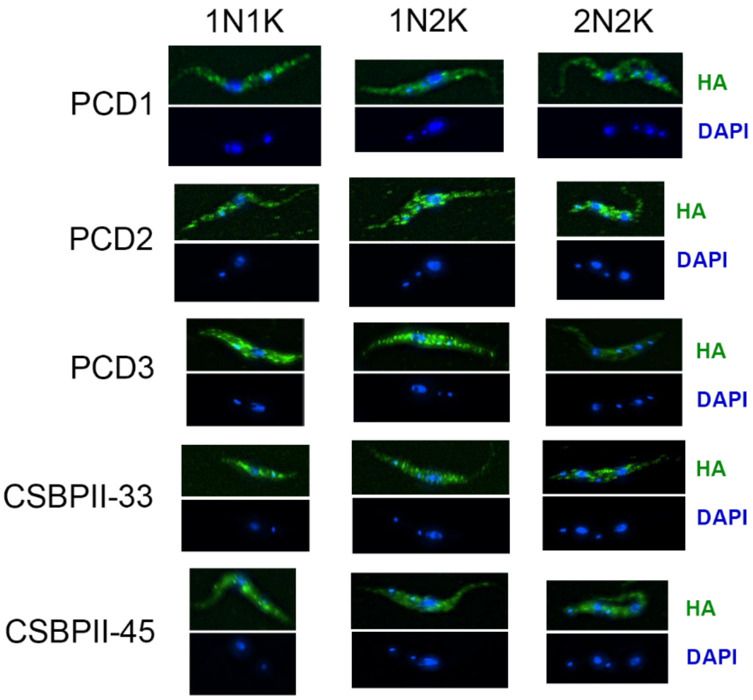

Figure S6. Localisation of cell cycle regulated PSP1-C domain containing proteins does not alter over the cell cycle. HA tagging endogenous tagging and immunofluorescence microscopy revealed the proteins have punctate localisation within the cytosol. No change in localisation occurred over the cell cycle, as judged by examining images with differing nucleus and kinetoplast counts.

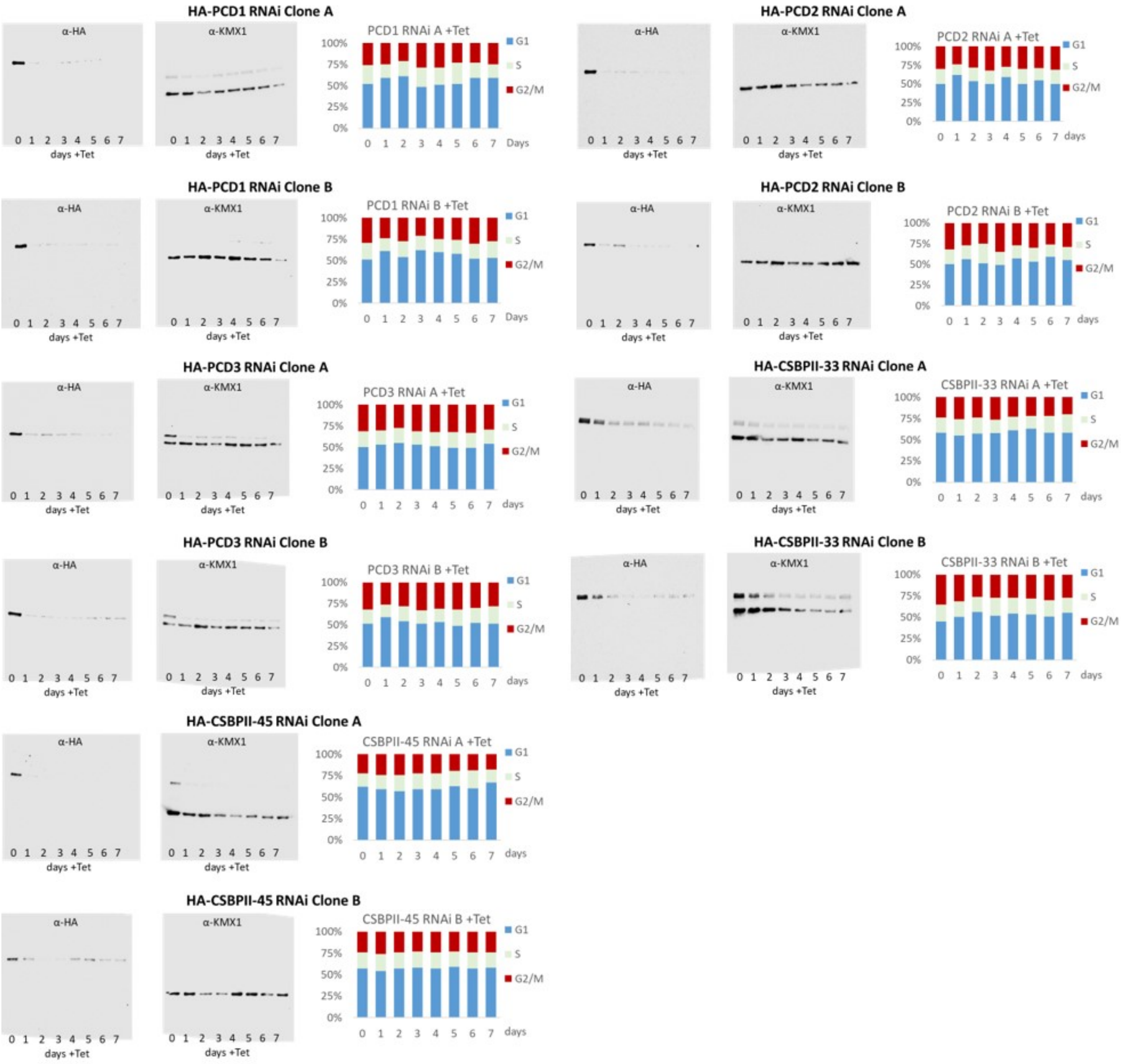

Figure S7. Tetracycline inducible RNAi of HA-tagged PCD proteins. Aliquots of cells from the respective RNAi time course were subjected to Western blotting and flow cytometry. Western blots with anti-HA confirmed efficient knockdown of the HA-tagged proteins, with an anti-KMX-1 (tubulin) used a loading control. Flow cytometry of PI-stained cells revealed that the proportion of cells in different cell cycle time points was unchanged.

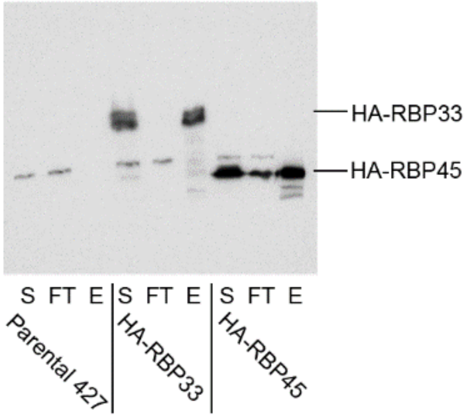

Figure S8. Immunoprecipitation of the T. brucei CSBPII complex proteins. IP of HA-*Tb*CSBPII-45, HA-*Tb*CSBPII-45, and the parental 427 cells with anti-HA beads, subjected to anti-HA western blotting. HA-tagged proteins can be observed in the eluent. S – starting material; FT – flow through; E – eluent.

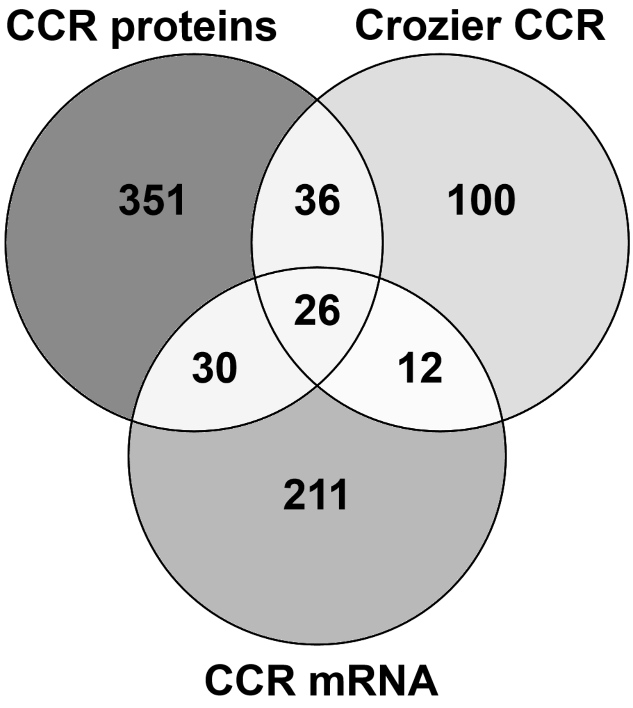

Figure S9. Venn diagram of the overlap of the two CCR proteomes and the CCR transcriptome. CCR proteins – 443 proteins identified in the present study; Crozier CCR – 174/384 CCR proteins reported by Crozier *et al.* were quantified at all six time points; CCR mRNA – 279/528 CCR transcripts reported by Archer *et al.* were quantified at all six time points.
